# Fungal hacking of the plant sex-determination pathway via interference with *AGL24* in *Silene latifolia*

**DOI:** 10.1101/2023.08.28.555088

**Authors:** Naoko Fujita, Michael E. Hood, Yuka Komoda, Takashi Akagi

**Affiliations:** Graduate School of Environmental and Life Science, Okayama University, Okayama, Japan; Amherst College, Amherst, Massachusetts, USA; Department of Sustainable Agriculture, College of Agriculture, Food and Environment Sciences, Rakuno Gakuen University, Hokkaido, Japan

## Abstract

Plants have evolved lineage-specific sex-determination systems that is determined not only by genetic factors, but also the surrounding environmental conditions, including interactions with pathogens. *Silene latifolia* is a model dioecious plant whose sexuality is genetically regulated by X/Y chromosomes; however, anther smut fungus mimics the plant Y chromosome and forcibly converts female plants to male. Here, transcriptome analyses of healthy or fungus-infected *S. latifolia* inflorescence meristems suggested that an orthologue of *AGL24* (*SlAGL24*), a flowering activator, is a key factor in sex conversion *via* fungus infection. Overexpression of *SlAGL24* in *Arabidopsis thaliana* suppressed stamen development, whereas knock-down of *SlAGL24* in *S. latifolia* converted males into hermaphrodites. Furthermore, *SlAGL24* expression affected sexual dimorphisms in *S. latifolia*. Our results propose an adaptive scenario wherein the anther smut fungus targets *SlAGL24*, as a master regulator connecting the fungal signal to sex determination, to confer male and potentially male-beneficial traits, effectively transmitting its teliospores.

## MAIN TEXT

Separated sexuality is one of the main strategies to maintain genetic diversity within a species, both in animals and plants. Plants have evolved lineage-specific chromosomal sex-determination systems, or separated male and female individuals (dioecy), from the ancestral functional hermaphroditism, potentially *via* various intermediate sexual systems^1^. Although dioecy is often viewed as an irreversible state, a so-called ‘dead-end’ in reproductive biology^2^, some plants have exhibited escape from dioecy to ancestral states such as monoecy^3^ or hermaphroditism^4, 5, 6, 7^. These flexible transitions have potentially served as an adaptive strategy to ensure a balance between genetic diversity and stable reproduction in immobile plants. This situation may also require substantial adaptive changes in organisms that live symbiotically (or parasitically) with plants, so that their life cycles adjust to the plant mating system. Pathogens have evolved various strategies to manipulate hosts for their own benefit, including sex change or “hacking” of the host reproductive systems. In insects, several endosymbiotic bacteria, such as *Wolbachia, Spiroplasma,* and *Rickettsia*, can affect the sex or sex ratio^8, 9^, ^10, 11^. These bacteria are transmitted exclusively through female hosts and have evolved the ability to bias populations towards females by turning males into females (feminization) or eliminating males (male killing), and to trigger clonal reproduction (parthenogenesis).

*Silene latifolia* (the family Caryophyllaceae) is a dioecious plant with a heterogametic male (XY) sex-determination system. It has long been used as a model plant for research on sex chromosome evolution, ever since sex chromosomes were first identified in angiosperms in 1923^12^. Consistent with the representative framework of the “two-mutation” model^14, 15^, two Y-chromosome-localized sex-determining genes, encoding male-promoting (M) and female-suppressing (SuF) factors, are thought to be responsible for the expression of dioecy^16, 17^. Recently, a homolog of Arabidopsis *CLAVATA3* was isolated as a candidate gene for SuF based on Y-deletion mapping of hermaphrodite mutants^18^. Infection by the anther smut fungus (*Microbotryum lychnidis-dioicae* (DC.) G. Demi & Oberw.) can make *S. latifolia* female (2A+XX) plants produce hermaphrodite flowers, despite lacking the Y chromosome (Figure 1b,e). This flower sex conversion caused by the anther smut fungus caused Carl Linnaeus to wrongly define the sex-converted hermaphrodites as a different species^19^. The anther smut fungus is an obligate biotrophic plant pathogen that forms a large number of infectious teliospores^20, 21^. Among the lineages of smut fungi, *M. lychnidis-dioicae* has a specific ability to produce teliospores by replacing *S. latifolia* pollen, thereby ensuring their transmission by insect pollinators. Theoretical frameworks assume that, to increase the chance of transmission, anther smut fungus has evolved a strategy to force females to produce stamens (or anthers) in dioecious *S. latifolia*^22^. Here, we explored the mechanism by which the anther smut fungus promotes male functions independently of Y-chromosome factors, and focused on the male-beneficial traits (or sexually antagonistic traits) that accompany this conversion.

**Figure 1.**
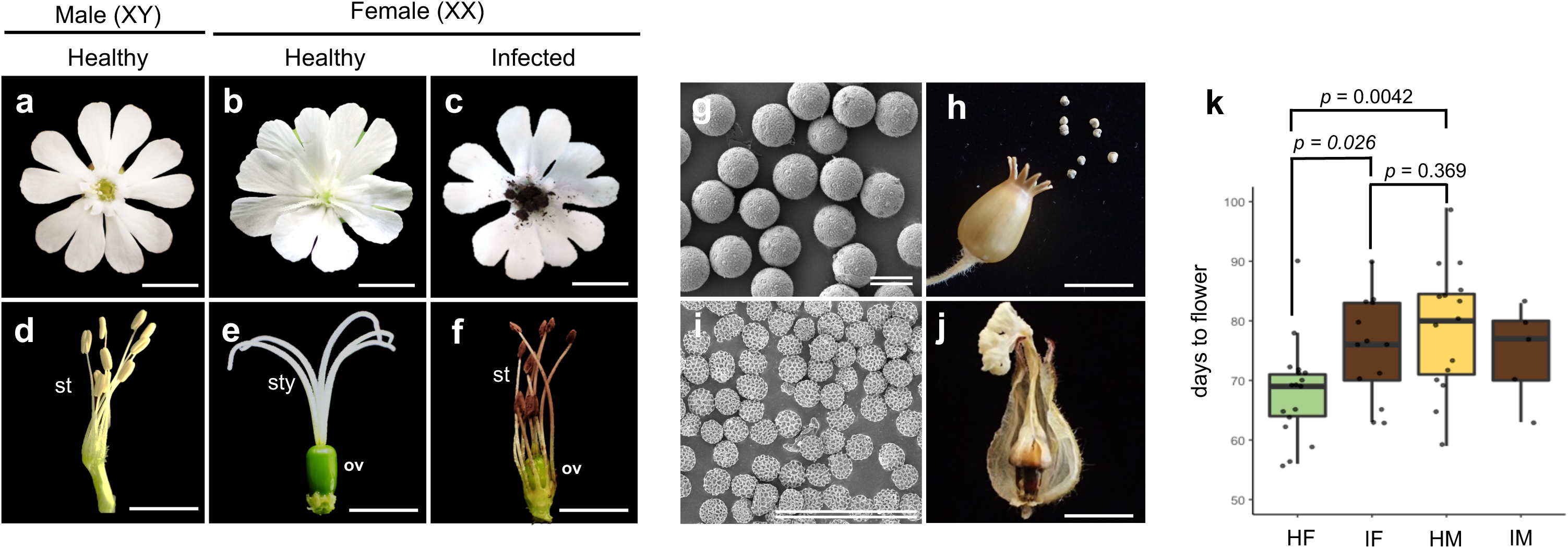
Characterization of the masculinized female flower infected with the anther-smut fungus. **a and d**, Genetically male (XY) flower with stamens (st) producing yellowish pollens in the anthers. **b and e**, Genetically female (XX) flower with functional gynoecium including ovary (ov) and five styles (sty). **c and f**, Masculinized female (XX) flower infected with the anther-smut fungus. Dissected flowers removing sepals and petals were shown in **d**-**f**. In the anther-smut fungus infected female (**c** and **f**), gynoecium development is severely arrested, while stamens can develop as in genetically male flowers. However, the pollens are replaced with the blown-purple fungal spores. **g,** Scanning electron micrograph of normal pollen grains in genetically male. **h,** Mature ovary resulted from pollination with normal pollen grains produce seeds. **i,** Scanning electron micrograph of fungal teliospores collected from the infected masculinized female flower, showing no ability to produce normal pollens. **j,** Withered ovary of the healthy (non-infected) female flower after pollination with the fungal spores from the masculinized female anthers. Bars = 1 cm (single line), 30 µm (double line). **k**, Days to flower after sowing in healthy female (HF), the infected masculinized female (IF), healthy male (HM), and infected male (IM). Genetically male (HM) and the masculinized infected female (IF) exhibited substantial delay in flowering time in comparison to healthy female (HF) (*p =* 0.0042, 0.026, Two-sided Student’s t-test with *N* >10 each phenotype group), while there is no significant difference in flowering time between IF and HM (*p* = 0.369, Two-sided Studen’s t-test with *N* > 10 each phenotype group).

We experimentally classified *S. latifolia* individuals into four categories according to their genotype and anther smut infection status; i.e., healthy female (HF), infected female (IF) (visually hermaphroditic with short pistil, or incomplete male, see Extended data Figure S1), healthy male (HM), and infected male (IM) (no sex conversion). The male and female functions of this species are thought to be independent under the two-mutation model^14, 16^, suggesting that the anther smut fungus not only promotes androecium development, but also suppresses gynoecium suppression. Our scanning electron microscope (SEM) observations demonstrated that IF anthers produced only fungal teliospores (Figure 1i) that were not recognized as pollen during pollination (Figure 1j). Considering the fact that IM anthers produced both functional pollen and fungal teliospores, the anther smut fungus was unable to promote perfect androecium development. Although the IF ovary partially matured even without pollination, it was filled with fungal spores (Extended data Figure S2). Importantly, anther smut infection complemented some sexual dimorphism traits. In *S. latifolia*, representative sexual dimorphism characters include flower number^23, 24^, flowering time^25, 26^, and flower size^24^. In our study, not only genetically male (or both HM and IM) but also masculinized infected female (or IF) plants exhibited substantial delays in flowering time compared with that of healthy females (HF) (*p* < 0.05, Student’s t-test) (Figure 1k). Infection by the anther smut fungus reduced the number of flowers in genetically male (IM) and female plants (IF), despite the higher number of flowers in male (HM) than in female (HF) plants (Extended data Figure S3).

To identify genes involved in the physiological changes during sex conversion caused by the anther smut fungus infection, we performed micro-dissecting RNA-seq analyses using early inflorescence meristems that were sampled before the stamen primordia became visible (Figure 2a–c). We searched for genes showing clear differences in their expression patterns between HF (visually female) and the other three visually male (or hermaphrodite) plants, i.e., IF, HM, and IM, hypothesizing that sex conversion (development of an androecium) *via* anther smut infection shares regulatory pathways with those functioning in genetic males. Hierarchical clustering of the pre-selected differentially expressed genes between HF and IF, HM, or IM (FDR<0.01 with DESeq2), revealed two candidate clusters, where IF/HM/IM synchronously showed upregulation (cluster 8, Figure 2e, Supplementary Table S1) or downregulation (cluster 4, Figure 2f, Supplementary Table S1) against HF. Cluster 8 contained many genes annotated with “Reproduction” (Figure 2g, Supplementary Table S1). They included, for example, *FRIGIDA-like proteins* (*FRL*s) and *GIGANTEA* (*GI*), which are known to be involved in the control of flowering time^27, 28, 29^. Cluster 4 contained genes involved in reproductive transition in the meristem, including *NUCLEOSTEMIN-LIKE 1* (*NSN1*), *AGAMOUS-LIKE 24* (*AGL24*), and *SUPPRESSOR OF OVEREXPRESSION OF CO 1* (*SOC1*) ^30, 31, 32, 33^ (Supplementary Table S1). Among them, *AGL24* encodes a repressor of class B, C, and E genes, all of which show fluctuations in expression that potentially affect both gynoecium and androecium development^33^. Importantly, *SHORT VEGETATIVE PHASE* (*SVP*), a close paralog of *AGL24,* acts as its antagonist, and is known to be involved in androecium development during sex differentiation in persimmon (genus *Diospyros*). *SVP* is positively regulated by the transcription factor MeGI, and SVP regulates the expression of *SOC1*^34^. Furthermore, the expression pattern of *AGL24* in *S. latifolia* might well be explained by the pleiotropic functions of *Arabidopsis AGL24*, which represses floral homeotic genes that control petal stamen and carpel identity^33, 35^, in addition to promoting flowering time^30,31^.

**Figure 2.**
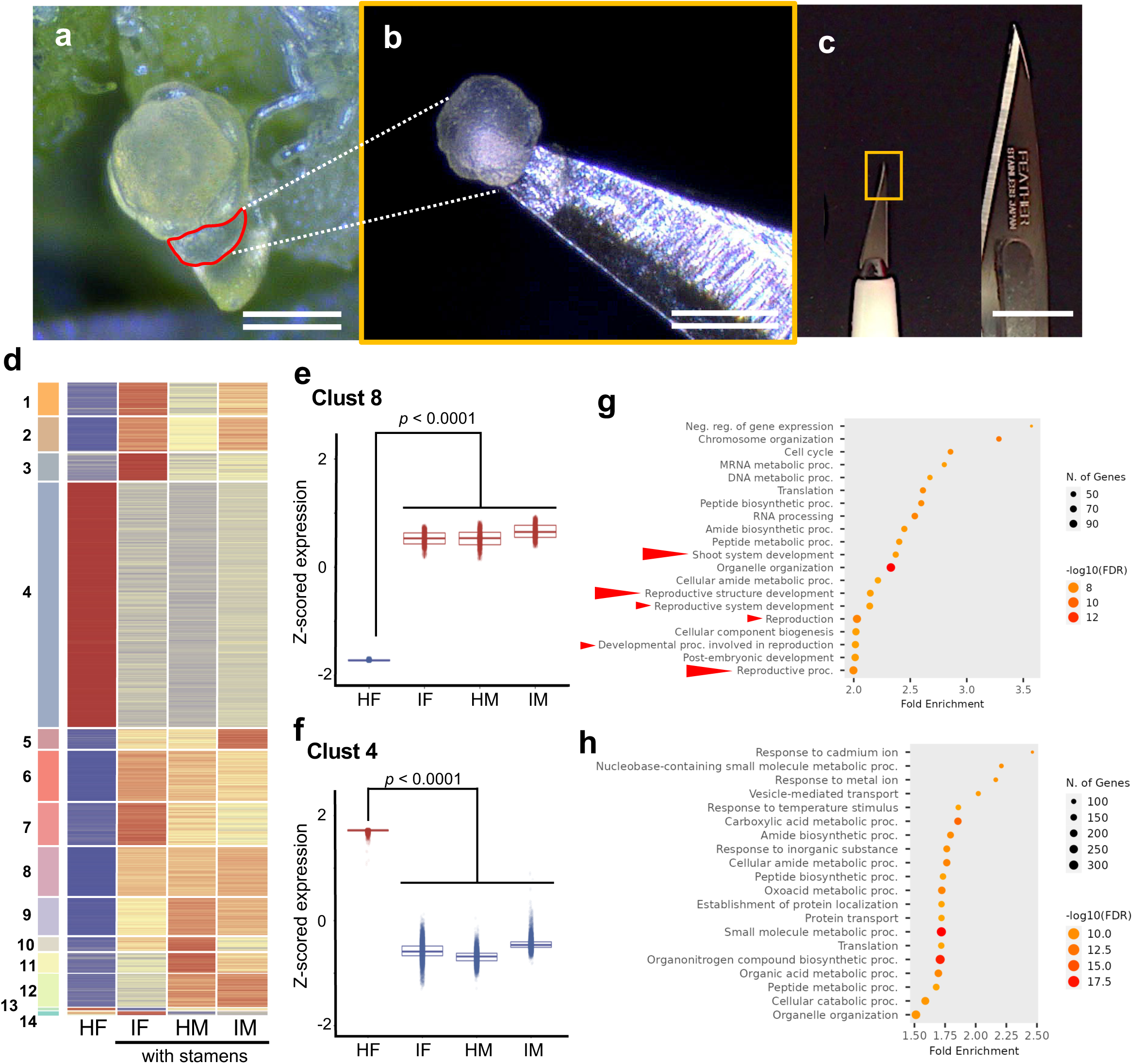
Micro-dissecting RNA-seq analyses for identification of the genes consistent with the sex expression. **a-c,** Micro-dissecting of early inflorescence meristems before the stamen primordia are visible. **a,** A representative inflorescence meristem used for this study is shown with red line. **b,** An inflorescence meristem dissected with the ophthalmology blade shown in **c,** compared with a normal lab-use blade. A tip of the ophthalmology blade shown with the yellow box corresponds the whole panel (**b**). Bars = 500 µm (single line), 100 µm (double line). **d,** Hierarchical clustered heatmap of differentially expressed genes between HM (healthy male) and HF (healthy female) or HF and IF (the infected female). **e,** Integrated expression pattern of Clust 8, constituted of the genes upregulated in male (IF, HM, and IM) which promoted stamens development. **f,** Integrated expression pattern of Clust 4, constituted of the genes downregulated in male (IF, HM, and IM). **g and h,** GO terms enriched in Clust 8 (**g**) and Clust 4 (**h**). Arrow heads indicated the terms related to physiological changes in the inflorescence meristem.

To characterize the functions of *S. latifolia AGL24* and its paralog *SVP* (namely, *SlAGL24 and SlSVP*, respectively), we transformed *Arabidopsis thaliana* plants with *SlAGL24* and *SlSVP* driven by the constitutive CaMV35S promoter. The transformants harbouring *SlAGL24* and those harbouring *SlSVP* exhibited suppressed growth of stamens and petals (Figure 3b–e), suggesting that *SlAGL24* and *SlSVP* play similar roles in repressing class B floral homeotic genes in the ABC model. The deficient anthers produced pollen with reduced viability (Figure 3g,h). The transformants harbouring *SlAGL24* showed precocious flowering, while those harbouring *SlSVP* showed a delayed flowering time (Figure 3i,j), consistent with the potentially antagonistic functions of AGL24 and SVP reported in other studies^30, 31, 36^. These observations in *SlAGL24*-overexpressing transgenic lines explained the phenotypic changes in IF (or converted to male) and males (HM and IM) that exhibited delayed flowering (Figure 1k) and androecium development (Figure 1a,f), accompanied by downregulation of *SlAGL24* (Figure 2f).

**Figure 3.**
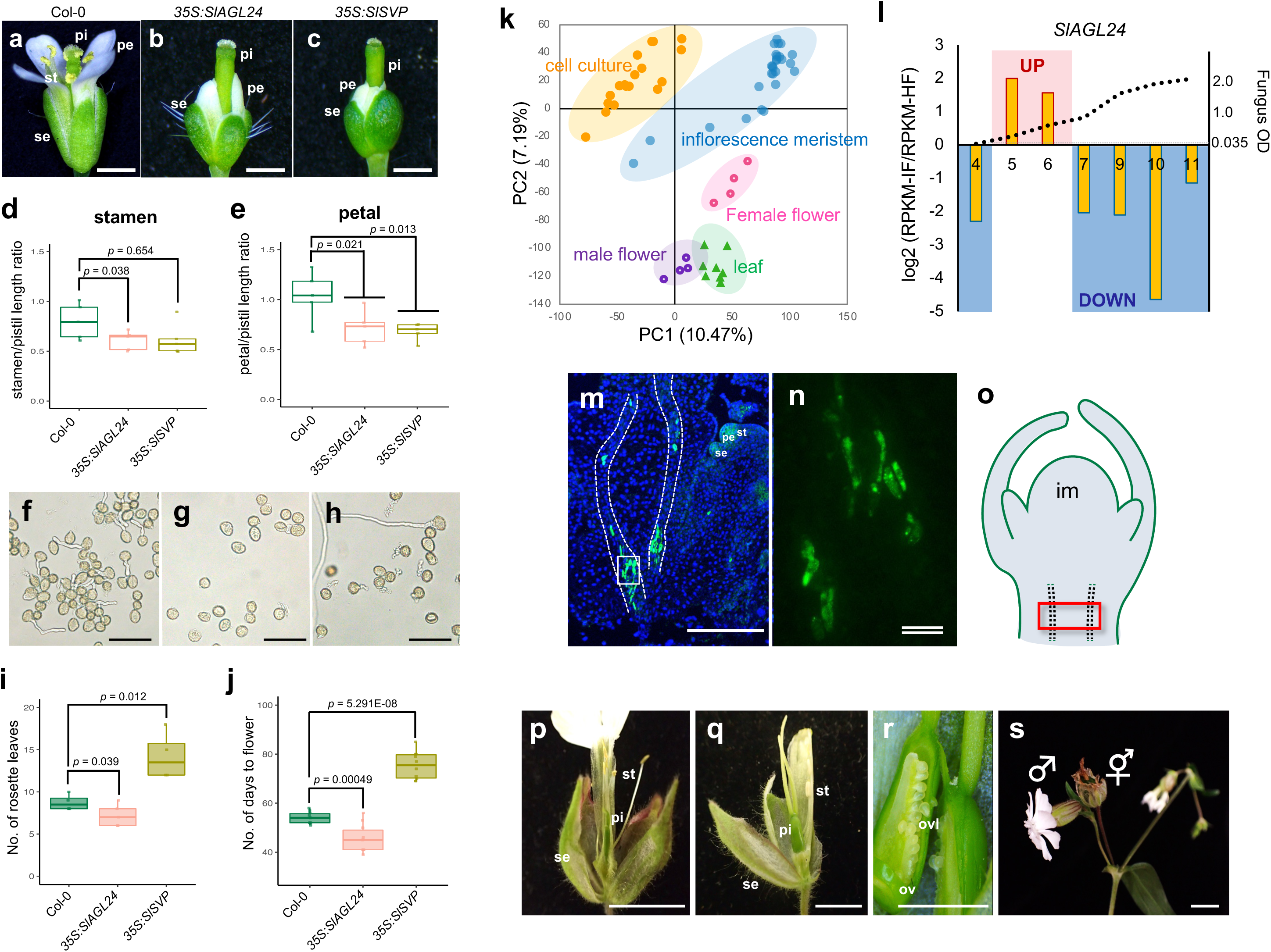
Pleiotropic functions of *SlAGL24* for sex determination. **a-j,** Constitutive expression of *SlAGL24* and *SlSVP* in *A. thaliana* plants. Representative flower phenotype of wild-type Col-0 (**a**), *35S:SlAGL24* (**b**), *35S:SlSVP* (**c**). st: stamen, pi: pistil, pe: petal, se: sepal. Bars = 1 cm. The *35S:SlAGL24* and *35S:SlSVP Arabidopsis* lines exhibited consistently suppressed stamens and petals development, while the stamen suppression effect was slightly less in *35S:SlSVP* than in *35S:SlAGL24* (**d-e**, *p* = 0.038 and 0.654 for stamens/pistil length ratio, 0.021 and 0.013 for petal/pistil length ratio, respectively. One-sided Student’s *t*-test with *N* = 5). Pollen tube growth (**f**: WT, **g**: *35S:AGL24*, **h**: *35S:SlSVP*) was severely arrested in *35S:SlAGL24* (**g**) and mildly suppressed in *35S:SlSVP* (**h**). Flowering time was examined with the number of rosette leaves (before bolting) (**i**) and the days to flower after sowing (**j**). *SlAGL24* induced precocious flowering (*p* = 0.039 for rosette leaves nos, and *p* = 0.00049 for days to flower), while *SlSVP* oppositely delayed flowering (p = 0.012 for rosette leaves nos, and *p* = 5.29E-08 for days to flower). **k,** Principal component analysis of the gene expression profiles in suspension cell cultures (*orange*), the inflorescence meristem (*blue*), female and male mature flowers (*pink* and *purple*, respectively), and leaves (*green*). **l,** Temporal *SlAGL24* expression fluctuation in the suspension cell cultures infected with the anther-smut fungus. Growth curve of the fungus (OD_600_) was shown with the dotted line. *N* = 2 for each measuring points. **m-o,** *in situ* localization of *SlAGL24* in the shoot apex of the infected-female hybridized with *SlAGL24* antisense probe, detecting strong signals shaping outer contour of the fungal cells in vascular bundle (dashed lines) under the shoot apex (**m**). se: sepal, pe: petal, st: stamen. An enlarged image of the white box in (**m**) exhibited fungus-shape-like *SlAGL24* signals (**n**). An illustration of the shoot apex (**o**). The position of the vascular bundle observed in (**m**) was given with the red box. im: inflorescence meristem. Bars = 100 µm (single line), 10 µm (double line). **p-s,** Hermaphroditic phenotype induced by *SlAGL24*-ALSV virus induced silencing (VIGS) in genetically male *S. latifolia* plants (**p** and **q**). A longitudinal section of the ovary in (**q**) contained functional ovules (**r**) as in healthy female flowers. A maturing ovary of hermaphrodite flower formed a fruit producing normal seeds by self-fertilization (**s**). st: stamen, pi: pistil, se: sepal, ov: ovary, ovl: ovules. Bar = 1 cm

A principal component analysis (PCA) was conducted using the transcriptomic data obtained from developing male and female flowers (Figure 1a–c), leaves, early inflorescence meristem (Figure 2a–c), and suspension-cultured cells derived from leaves (see Methods section). The PCA results showed that the expression dynamics in early inflorescence meristems were similar to those in suspension-cultured cells (Figure 3k), with synchronous expression patterns of *AGL24* during anther smut fungus infection, as shown later. To detect fluctuations in the expression of *AGL24* over time during infection with the anther smut fungus, we monitored its transcript levels in an *in vitro* suspension cell culture directly inoculated with fungal sporidia. The expression bias between IF (infected) and HF (healthy) cells was negatively correlated with fungal growth in IF culture (Figure 3l, Pearson’s *r* = −0.51 for the expression bias of infected/healthy *vs.* fungus abundance). Importantly, in the initial fungus growth stage (day 5–6), *SlAGL24* expression was temporarily upregulated (infected/healthy >1.5), while in the later fungus growing stage (day 7–11), *SlAGL24* was downregulated (infected/healthy <-2.0). A similar expression pattern was observed in the male suspension cultured cells (Extended data Figure S5). Furthermore, in an *in situ* RNA hybridization analysis with the *SlAGL24* sequence as a probe, there were strong signals in the shape of the outer contour of the fungal cells in the vascular bundle under the shoot apex in IF (Figure 3m–o). The genome of anther smut fungus^37^ does not contain any *SlAGL24*-like sequences (< 1e^-2^ for the threshold of *p*-values in blastn), suggesting that the *SlAGL24* signal was directly activated in *S. latifolia* cells by anther smut fungus infection. These results suggest that, although *SlAGL24* expression can be controlled by the anther smut fungus, the regulatory pattern is spatiotemporally differentiated, rather than simply downregulated, as observed in the initial development stage of flower meristems.

Knocking-down of *SlAGL24* in male *S. latifolia* plants by apple latent spherical virus (ALSV)-mediated virus-induced gene silencing (VIGS)^38^ resulted in the production of hermaphrodite flowers (Figure 3r–u, Supplementary Table S1). These feminized hermaphrodites produced self-pollinated progeny that segregated into male and female individuals with an equal ratio (Supplementary Table S2), with no hermaphrodite offspring. In *S. latifolia*, the Y chromosome cannot be transmitted through the egg^39^. Hence, the expected sex ratio would be 1:1 female:male in the progeny derived from hermaphroditic male (XY) self-pollination, consistent with our results. The fact that the hermaphroditic phenotype was not inherited by the progeny confirmed the non-genetic silencing of *SlAGL24* by VIGS.

On the basis of our results, we speculate that *SlAGL24* is the hub gene through which the anther smut fungus hacks the sex-determination pathway in *S. latifolia*. *SlAGL24* can contribute pleiotropically to androecium development and gynoecium suppression, and this is achieved by the modification of its expression by anther smut fungus infection in genetically female plants, as shown in Figure 4. In our results, *SlAGL24* was both upregulated and downregulated during anther smut fungus infection, although further research is required to explore its spatiotemporal expression patterns in more detail. Reciprocal effects of *AGL24* (or *AGL24/SVP* family) on androecium and gynoecium development (or class B/C floral homeotic genes) have been proposed in *Arabidopsis*^33^, where the direction of the effect strongly depends on the floral developmental stage. A protein related to the genetic sex-determination pathway in *S. latifolia*, functioning of the suppressor of feminization (SuF, or SlCLV3 as a candidate), is thought to precede the hypothetical male promoting factor (M), in *S. latifolia* male flowers^40^. Induction of stamens in female plants infected with anther smut fungus occurs synchronously with the functioning of this genetic pathway^20^. Our findings that downregulation of *SlAGL24* expression occurs in both genetically male and anther smut fungus-infected female plants, at least during flower development stages 1 to 4, suggest that *SlAGL24* is under the control of the M factor for androecium development. However, more direct evidence is required to confirm whether and how SuF (or SlCLV3) regulates *SlAGL24*, since we have not detected the point at which *SlAGL24* is upregulated in the (defective) gynoecium primordia of genetically male plants. The fluctuations in *SlAGL24* expression regulated by anther smut fungus infection affect not only sex expression, but also the sexual dimorphism character of flowering time. This result also implies that the Y-chromosome-localized sex-determining genes have pleiotropic functions to directly regulate sexual dimorphism *via* the regulation of *SlAGL24* expression, as previously suggested in persimmon and kiwifruit^41, 42^. Thus, the ability of anther smut fungus to target *SlAGL24* might be an adaptive and reasonable strategy to cause female *S. latifolia* to mimic a male plant, with potentially beneficial male traits to effectively transmit pollen (or teliospores in infected females).

**Figure 4.**
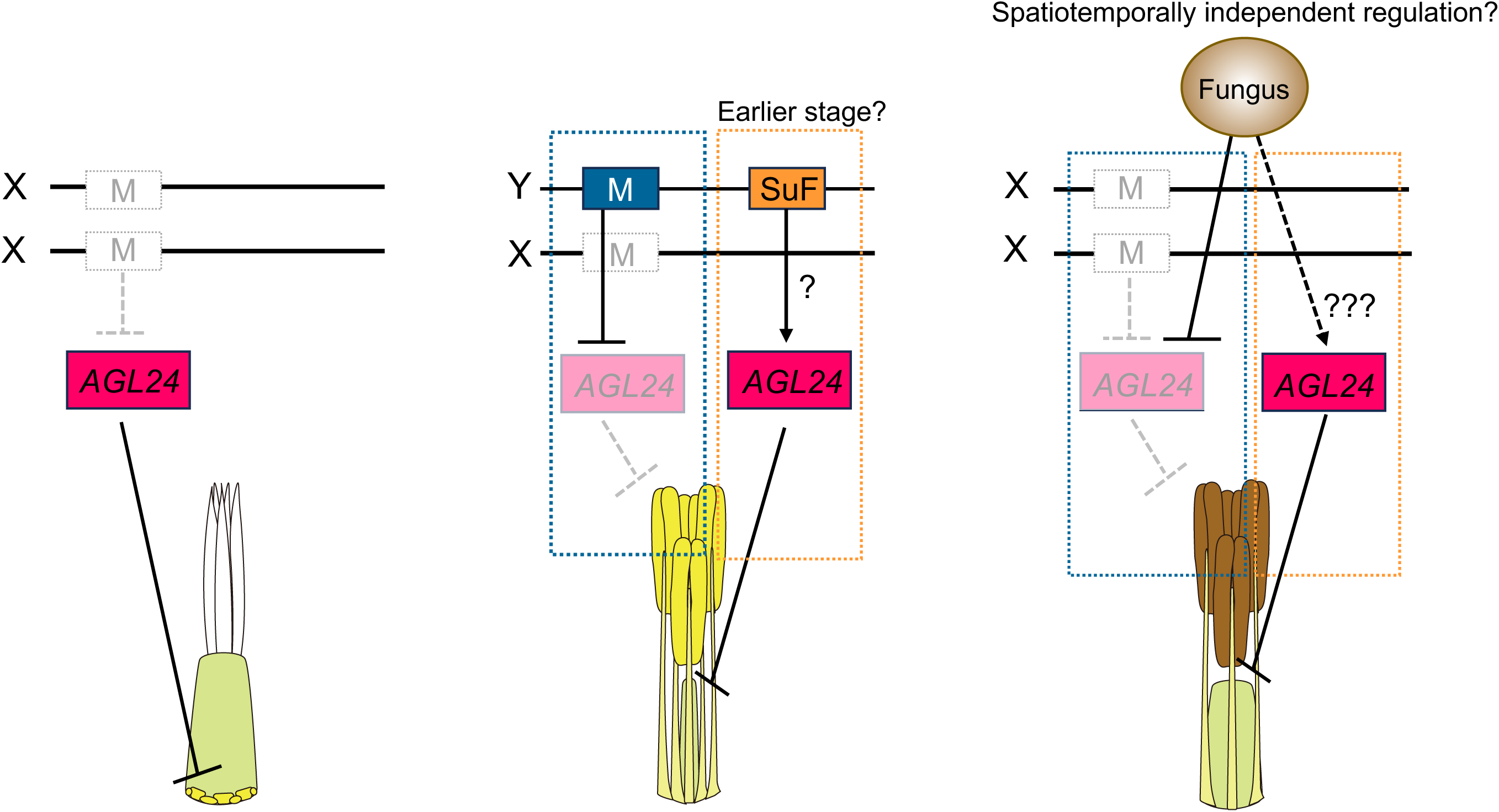
Model for fungal hacking of the plant sex-determination pathway via interference of *AGL24*. Spatiotemporally regulation of *AGL24* the sex-determination pathway in *S. latifolia* plants. In genetically female (XX) flowers (*left*), upregulation of *AGL24* arrests the development of stamens which is under the control of a putative male factor (M). In genetically male (XY) flowers (*middle*), downregulation of *AGL24* promotes the development of stamens through the function of M on the Y chromosome. Spatiotemporally downregulation of *AGL24* promoted the gynoecium implied that AGL24 is regulated by not only M but also female suppressing (SuF) factor. In the anther-smut fungus infected female (*right*), the fungus represses the expression of *AGL24* by acting as M factor. A partial repression of the gynoecium development may be caused by spatiotemporally independent regulation of *AGL24* by the fungus.

## METHODS

### Plant and fungus materials

*Silene latifolia* seeds used in this study were the inbred lines from the previous study^38^. The seeds were surface sterilized and sowed on Murashige and Skoog (MS) media in a growth chamber under conditions of 23 °C with a daylength of 16 hr. Two weeks after sowing, at cotyledon stage, *Microbotryum lychnidis-dioicae* Lamole strains^43^ were inoculated as follows. Haploid sporidium of Lamole A1 and A2 strains grown on potato dextrose agar were suspended in distilled water at 4×10^6^ cells/mL. After incubation in dark at 16°C overnight, equal volume of the Lamole A1 and A2 solution were mixed and applied onto the shoot apical meristems. For control, distilled water was applied instead of the fungal sporidium. The plants were grown on MS media for a month (four to six leaves) and transplanted to soil (Kumiai Nippi Engei Baido No.1, Nihon Hiryo Co., Ltd., Tokyo, Japan) in a pot (Φ 9 cm x 7.5 cm).

All plants used for the measurement experiments were grown in the same growth camber under constant conditions (23°C, 16L/8D). Flowering time was measured by counting the number of days from sowing to the first flower opened. Flower numbers determined by the number of branches were counted after two weeks from the first flower.

### Scanning electron microscopy

The anthers of healthy male and the infected female flowers were fixed with 2% paraformaldehyde (PFA) and 2% glutaraldehyde (GA) in 0.05 M cacodylate buffer pH 7.4 at 4°C overnight. The samples were additionally fixed with 1% tannic acid in 0.05 M cacodylate buffer pH7.4 at 4°C for 4 h. After the fixation the samples were washed four times with 0.05 M cacodylate buffer for 30 min each, and postfixed with 2% osmium tetroxide (OsO4) in 0.05 M cacodylate buffer at 4°C overnight. The samples were dehydrated in graded ethanol solutions (50%, 70%, 90%, anhydrous), subsequently the samples were continuously dehydrated in anhydrous ethanol at room temperature overnight. The samples were substituted into tert-butyl alcohol to freeze at 4°C. The frozen samples were vacuum dried. After drying, the samples were coated with thin layer (30 nm) of osmium by using an osmium plasma coater (NL-OPC80A; Nippon Laser & Electronics Laboratory, Nagoya, Japan). The samples were observed by a scanning electron microscope (JSM-7500F; JEOL Ltd., Tokyo, Japan) at an acceleration voltage of 3.0 kV.

### Cell culture and fungus inoculation

*S. latifolia* leaves were sterilized for 10 minutes with 1% sodium hypochlorite (NaOcl) and 1% Tween-20 in distilled water, followed by three times for 10 mins in sterile distilled water. The explants were cut into 5 mm in rectangles on filter paper, and placed on callus-inducing medium containing 1x MS (M5519-50L; Sigma-Aldrich, St. Louis, MO, US), 3% sucrose, 1 mg/L 2.4-D, 0.1 mg/L BA, 0.25% gellan gum (FUJIFILM Wako Pure Chemical Corp., Osaka, Japan) pH 5.7, which were incubated at 23°C under dark conditions. Calli which became visible after one to two weeks were maintained by refreshing the media every four weeks.

For generation of suspension cell cultures, the calli chopped with a tweezers were suspended in liquid media with the same composition as the callus-inducing medium without adding gellan gum. The culture cells were kept on a rotary shaker at 105 rpm at 23°C under dark conditions. For fungal inoculation, 10 µl of the fungal sporidia at concentration of 2×10^6^ cells/mL which equally contained Lamole A1 and A2 fungal strains were added into the suspension cell cultures.

### Transcriptome analysis

Inflorescence meristems at early stages (stage 1-4^40^) were collected under stereomicroscope Olympus SZ (Evident Co., Tokyo, Japan) with an ophthalmic surgical knife P715 (Feather Safety Razor CO., Ltd, Osaka, Japan). Suspension culture cells were collected by centrifugation. Total RNA was isolated using Favorgen Plant Total RNA Mini Kit (Favorgen Biotech Corp., Ping-Tung, Taiwan). Extracted total RNA were processed in preparation for Illumina Sequencing as follows. First, mRNA was purified using the Dynabeads mRNA purification kit (Life Technologies). Next, cDNA was synthesized via KAPA RNA HyperPrep kit (Roche), followed by a DNA cleanup step with AMPure XP beads (Beckman Coulter; AMPure:reaction, 0.8:1). The constructed libraries were sequenced on Illumina’s HiSeq 4000 (50-bp single-end reads) or HiSeqX (150-bp paired-end reads). Illumina sequencing was conducted at the Vincent J. Coates Genomics Sequencing Laboratory at UC Berkeley for Hiseq 4000 or at Macrogen for HiseqX. Raw sequencing reads were processed using Python scripts (https://github.com/Comai-Lab/allprep/blob/master/allprep-13.py) for preprocessing and demultiplexing of sequencing data. The obtained reads were mapped to male reference prepared by de novo assembly of coding sequences (CDSs) using Burrows – Wheeler aligner (BWA) (version 0.7.15, Li and Durbin 2009). Read counts per CDS were generated from the aligned SAM files using R script. PCA of mRNA expression was conducted using the genes with RPKM > 1, using prcomp in R. Differential expression between HF and IF/HM/IM was analyzed using DESeq2 package^44^. The DEGs were filtered according to RPKM and FDR values (RPKMC>C1, FDRC<C0.01). Clustering of the DEGs was conducted using the hclust function in R, and visualized via a heatmap using the ComplexHeatmap package. Putative functions of each gene were determined with a BLASTX search of the TAIR10 database (https://www.arabidopsis.org/index.jsp). A GO enrichment analysis was performed on the genes in the three networks using ShinyGO 0.77 (http://bioinformatics.sdstate.edu/go/).

### In situ hybridization

Buds were fixed in FAA (10% formalin, 5% acetic acid, 50% ethanol) on ice under vacuum for 4 hours. After replacing with freshly prepared FAA solution, the buds were kept on ice for overnight. The fixed tissues were washed three times with PBS for 10 mins, dehydrated in an ethanol series (25, 50, 75, 95, and 100%, each step for 20 mins on ice), and stored in 100% ethanol overnight at 4°C. The ethanol was replaced with fresh 100% ethanol. The tissue samples were immersed in 20% sucrose solution 24 h before sectioning. Tissue sections were prepared by Kawamoto’s film method^45^. An RNA probe for *SlAGl24* was prepared using a *AGL24*-specific primers (AGL24_111F, 5’- TCTTTGTGATGCCGATCTTG-3’ and AGL24_523R, 5’-ACATGGCCAACCTTTGTTTC-3’). The amplified fragment cloned into pGEM-T Easy vector (Promega K. K., Tokyo, Japan) was used to synthesize DIG-labeled sense and antisense RNA probes using DIG RNA labeling kit (SP6/T7) (Merck KGaA, Darmstadt, Germany). The DIG-labeled RNA probes were purified using ProbeQuant G-50 Micro Colums (Cytiva, Tokyo, Japan). *In situ* hybridization was performed based on the manual for a commercial kit ‘*In situ* Hybridization Reagents (ISHR)’ (Nippon Gene Co., Ltd., Tokyo, Japan). Briefly, the specimens were washed three times with PBS for 10 mins, treated with 5 µg/mL Proteinase K (New England Biolabs Inc., MA, US) for 5 mins, washed with PBS-Glycine buffer (2 mg/mL in PBS pH7.4) for 10 mins, and washed again two times with PBS for 3 mins. The tissue sections were acetylated using acetylation reagents ISHR3 and 4 (Nippon Gene Co., Ltd., Tokyo, Japan), washed with PBST (1% TritonX-100 in PBS) for 30 mins, and washed again three times with PBS for 5 mins. After pre-hybridization with the Hybridization buffer ISHR7 (Nippon Gene Co., Ltd., Tokyo, Japan) at room temperature for 2 h, the RNA probes (5 µg/mL) were hybridized at 55°C overnight. Anti-DIG-FITC (Merck KGaA, Darmstadt, Germany) was used for signal detection. The prepared specimens were stained with SlowFade Gold Antifade Mountant with DAPI (Invitrogen, MA, US) and were observed using a microscope Olympus BZ51 (Evident Co., Tokyo, Japan).

### Virus induced gene silencing

Virus induced gene silencing (VIGS) was performed as previously described^37^. Briefly, SlAGL24-ALSV vector containing 108 bp fragment of SlAGL24 (shown in Extended data Figure S6) was prepared by introducing the synthesized DNA fragment (Azenta Life Sciences, MA, US) to SalI-BamHI site of the ALSV vector^37^. The resulting SlAGL24-ALSV vector transformed into *Agrobacterium tumefaciens* EHA105 was use to inoculate into *Nicotiana benthamiana* plants by agro-infiltration. Viral RNAs were purified from the infected *N. benthamiana* plants and was precipitated onto gold particles. For viral inoculation, the cotyledons of *S. latifolia* seedlings in an emergence stage were bombarded at 40 psi by a particle bombardment using GDS-80 (Nepa Gene Co., Ltd., Chiba, Japan).

### Accession numbers

All Illumina sequencing data have been deposited in the DDBJ database: Short Read Archives (SRA) database (BioProject accession: PRJDB16382, Run ID SAMD00635100-00635152 and SAMD00636369-00636376 for Illumina sequencing reads).

## SUPPLEMENTARY INFORMATION

Extended Figure S1-S6

Supplementary Data S1-S2

## ACKNOWLEDGEMENT

We thank Dr. Luca Comai (Dept. Plant Biol and Genome Center University of California Davis, USA) and Dr. Deborah Charlesworth (Institute of Evolutionary Biology, University of Edinburgh) for discussion and comments for this study, Dr. Kyoko Watanabe (College of Agriculture, Tamagawa University) and Dr. Koichiro Ushijima (Graduate School of Environmental and Life Science, Okayama University) for supervision, Rika Uchida and Yuka Hon-iden for plant maintenance and technical assistance, Kojiro Matsumoto (Nepa Gene Co., Ltd.) for continuous supports of the gene gun GDS-80. This work was supported by Grant-in-Aid for JSPS Fellow JSPS Grant Number [JP18J40290 and JP23KJ1615] and Grant-in-Aid for Scientific Research (C) JSPS KAKENHI [JP20K06016 and JP23K05226] to N.F. and Grant-in-Aid for Transformative Research Areas (A) from JSPS [22H05172 and 22H05173] to T.A.

## AUTHOR CONTRIBUTION

N.F. conceived the study and designed experiments. N.F. and Y.K. conducted the experiments.

N.F. analyzed the data. N.F., T.A., and M.H. drafted the manuscript. All authors approved the manuscript.

## REFERENCES

1. Renner, S. S. The relative and absolute frequencies of angiosperm sexual system: dioecy, monoecy, gynodioecy, and an updated online database. Am J Bot. 101, 1588–1596 (2014)

2. Heilbuth, J. C. Lower species richness in dioecious clades. Am Nat. 156, 221–241 (2000)

3. Akagi, T., Henry, I. M., Kawai, T., Comai, L. & Tao, R. Epigenetic regulation of the sex determination gene MeGI in polyploid Persimmon. Plant Cell. 28, 2905–2915 (2016)

4. Wang, J. et al. Sequencing papaya X and Y^h^ chromosomes reveals molecular basis of incipient sex chromosome evolution. Proc. Natl. Acad. Sci. U.S.A 109, 13710–13715

5. VanBuren, R. et al. Origin and domestication of papaya T^h^ chromosome. Genome Res. 25, 524–533 (2015)

6. Massonnet, M. et al. The genetic basis of sex determination in grapes. Nat. Commun. 11, 1–12 (2020).

7. Masuda, K. et al. Reinvention of hermaphroditism via activation of a RADIALIS-like gene in hexaploid persimmon. Nat. Plants 8, 217–224 (2022).

8. Jiggins, F. M., Hurst, G.D. & Majerus M. E. Sex-ratio-distorting Wolbachia causes sex-role reversal in its butterfly host. Proc Biol Sci. 267, 69–73 (2000)

9. Zeh, D. W., Zeh, J.A. & Bonilla, M.M. Wolbachia, sex ration bias and apparent male killing in the harlequin beetle riding pseudoscorpion. Heredity 95, 41–49 (2005)

10. Hackett, K. J., Lynn, D. E., Williamson, D. L., Ginsberg, A. S. & Whitcomb, R. F. Cultivation of the Drosophia sex-ratio spiroplasma. Science 232, 1253–1255 (1986)

11. Kikuchi, Y. & Fukatsu, T. Rickettsia infection in natural leech populations. Microb Ecol 49, 265–271 (2005)

12. Blackburn K. B. Sex chromosomes in plants. Nature. 112, 687–688 (1923)

13. Winge, O. On sex chromosomes, sex determination and preponderance of female in some dioecious plants. C R Trav Lab Carlsberg. 15, 1–25 (1923)

14. Westergarrd、M. The mechanism of sex determination in dioecious plants. Adv. Genet. 9, 217–281 (1958)

15. Charlesworth B. and Charlesworth D. A model for the evolution of dioecy and gynodioecy. Am. Nat. 112, 975–997 (1978)

16. Westergaard, M. Structural changes of the Y-chromosomes in the offspring of polyploid Melandrium. Hereditas 32, 60–64 (1946)

17. Kazama, Y. et al. A new physical mapping approach refines the sex-determining gene positions on the Silene latifolia Y-chromosome. Sci Rep. 6, 18917 (2016)

18. Kazama, Y. et al. A CLAVATA3-like gene acts as a gynoecium suppression function in white campion. Mol Biol Evol 39, msac195 (2022) 10.1093/molbev/msac195

19. Linnaeus, C. Hortus Cliffortianus. Amsterdam (1737)

20. Uchida, W., Matsunaga, S., Sugiyama, R., Kazama, Y. & Kawano, S. Morphological development of anthers induced by the dimorphic smut fungus Microbotryum violaceum in female flowers of the dioecious plant Silene latifolia. Planta 218, 240–248 (2003)

21. Uchida, W., Matsunaga, S. & Kawano, S. Ultrastructural analysis of the behavior of the dimorphic fungus Microbotryum violaceum in fungus-induced anthers of female Silene latifolia flowers. Protoplasma. 226, 207–216 (2005)

22. Becker, H. G. Infection of species of Melandrium by Ustilago violaceae (Pers.) Fuckel and the transmission of the resultant disease. Annals of Botany 11, 333–348 (1947)

23. Laporte, M. M. & Delph, L. F. Sex-specific physiology and source-sink relations in the dioecious plant Silene latifolia. Oecologia 106, 63–72 (1996)

24. Steven, J. C., Delph, L. F., & Brodie, E. D. III Sexual dimorphism in the quantitative-genetic architecture of floral, leaf, and allocation traits in Silene latifolia. Evolution 61, 42–57 (2007)

25. Purrington, C.B. & Schmitt, J. Consequences of sexually dimorphic timing of emergence and flowering in Silene latifolia. J. Ecol. 86, 286–393 (1998)

26. Doust, J. L., O’Brien, G. & Doust, L. L. Effect of Density on secondary sex characteristics and sex ratio in Silene alba (Caryophyllaceae). Am. J. Bot. 74, 40–46 (1987)

27. Fowler S. et al. GIGANTEA: a circadian clock-controlled gene that regulates photoperiodic flowering in Arabidopsis and encodes a protein with several possible membrane-spanning domains. EMBO J. 18: 4679–4688 (1999)

28. Mizoguchi, T. et al. Distinct roles of GIGANTEA in promoting flowering and regulating circadian rhythms in Arabidopsis. Plant Cell 17, 2255–2270 (2005)

29. Wang, X., Gingrich D. K., Deng, Y. & Hong, Z. A nucleostemin-like GTPase required for normal apical and floral meristem development in Arabidopsis. Mol Biol Cell. 23, 1446–56 (2012)

30. Yu, H., Xu, Y., Tan E. L. & Kumar P. P. AGAMOUS-LIKE 24, a dosage-dependent mediator of the flowering signals. Proc. Natl. Acad. Sci. U.S.A. 99, 16336–16341 (2002)

31. Michaels, S. D. et al. AGL24 acts as a promoter of flowering in Arabidopsis and is positively regulated by vernalization. Plant J. 33, 867–874 (2003)

32. Liu, C. et al. Direct interaction of AGL24 and SOC1 integrates flowering signals in Arabidopsis. Development 135, 1481–1491 (2008)

33. Gregis V., Sessa A., Dorca-Fornell C., Kater M. M. The Arabidopsis floral meristem identity genes AP1, AGL24 and SVP directly repress class B and C floral homeotic genes. 2009 Plant J. 60: 626-637

34. Yang, H. W., Akagi, T., Kawakatsu, T. & Tao, R. Gene networks orchestrated by MeGI: a single-factor mechanism underlying sex determination in persimmon. Plant J. 98, 97–111 (2019)

35. Gregis, V., Sessa, A., Colombo, L. & Kater, M. M. AGL24, SHORT NEGETATIVE PHASE, and APETALA1 redundantly control AGAMOUS during early stages of flowering development in Arabidopsis. Plant Cell 18, 1373-1382 (2006)

36. Hartmann, U. et al. Molecular cloning of SVP: a negative regulator of the floral transition in Arabidopsis. Plant J. 21, 351–360 (2000)

37. Perlin, M. H. et al. Sex and parasites: genomic and transcriptomic analysis of Microbotryum lychnidis-dioicae, the biotrophic and plant-castrating anther smut fungus. BMC Genomics. 16, 461 (2015)

38. Fujita, N. et al. Development of the VIGS system in the dioecious plant Silene latifolia. Int. J. Mol. Sci. 20, 1031 (2019)

39. Janoušek, B., Grant, S. & Vyskot, B. Non-transmissibility of the Y chromosome through the female line in androhermaphrodite plants of Melandrium album. Heredity 80, 576–583 (1998).

40. Grant, S., Hunkirchen, B. & Saedler, H. Developmental differences between male and female flowers in the dioecious plant Silene latifolia. Plant J. 6, 471–480 (1994)

41. Akagi, T. & Charlesworth, D. Pleiotropic effects of sex-determining genes in the evolution of dioecy in two plant species. Proc. R. Soc. B.28620191805 (2019)

42. Akagi, T. et al. Recurrent neo-sex chromosome evolution in kiwifruit. Nat. Plants 9, 393–402 (2023)

43. Hood, M. E. Dimorphic mating-type chromosomes in the fungus Microbotryum violaceum. Genetics. 160, 457–461 (2002)

44. Love, M. I., Huber, W. & Anders, S. Moderated estimation of fold change and dispersion for RNA-seq data with DESeq2. Genome Biol. 15, 1–21 (2014).

45. Kawamoto, T. Kawamoto, K. Preparation of thin frozen sections from nonfixed and undecalcified hard tissues using Kawamoto’s film method (2012). Methods Mol. Biol. 2014, 1130, 149–164

