## Extended data for "Fungal hacking of the plant sex-determination pathway via interference with *AGL24* in *Silene latifolia*"

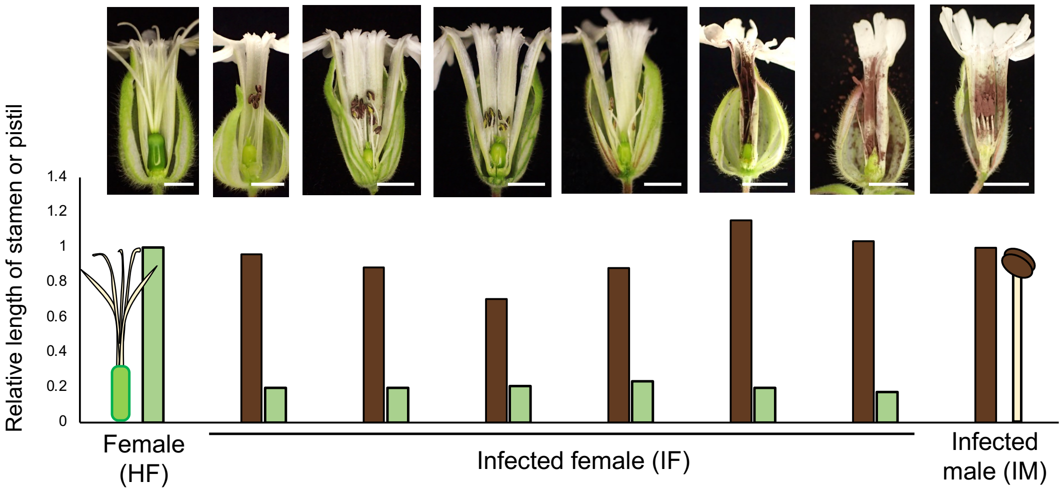


**Extended Data Fig.1: No correlation between stamen elongation and pistil suppression in the anther smut fungus infected female flowers**

Relative lengths of the suppressed pistil (*green*) and the induced stamen (*brown*) in the infected female (IF) compared with fully developed pistils in genetically female (HF, *left*) and stamens in the infected male (IM, *right*), respectively, are shown on the bar graph. Note that IM exhibited no sex change, with normal stamen development similar to a genetically male flower, while anthers are smutted. The stamen lengths were likely correlated to the number of fungal spores produced in the anthers. Each corresponding phenotype is shown in the top raw. Bars = 5 mm.

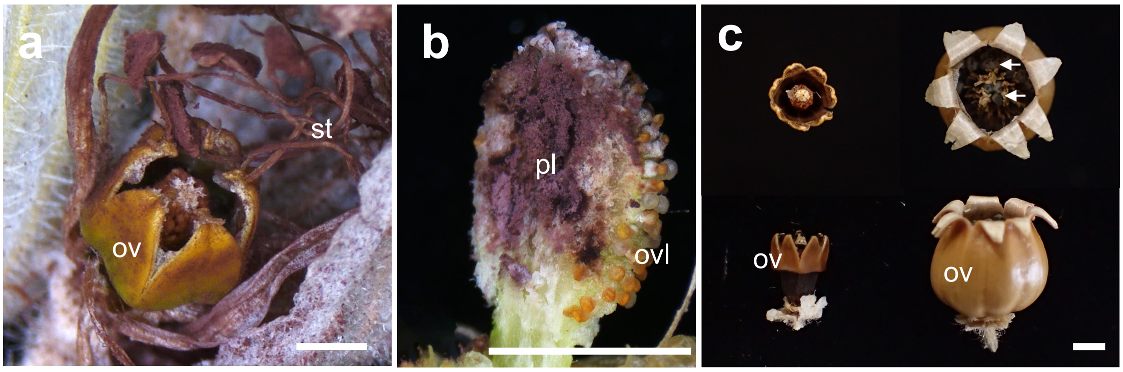


**Extended Data Fig.2: Smutted ovary of the anther smut fungus infected-female**

**a,** Smutted stamens (st) and partially matured ovary (ov) in the infected-female flower. For this self-pollination of female flower, fungal spores might be able to act as pollination stimulus, or viable pollen partially remained in the smutted anther **b,** whereas the placenta (pl) was filled with the fungal spores**.** Immature ovules (ovl) were observed in the ovary. **c,** Subsequently developed ovary of the anther smut fungus infected-female (*left*) at the stage of seed production in a control pollinated female flower (*right*). A number of seeds (arrow) were produced in the pollinated healthy female, whereas no seeds were produced in the anther smut fungus infected-female, showing that self-pollination does not occur with the fungal spores. Bars = 3 mm

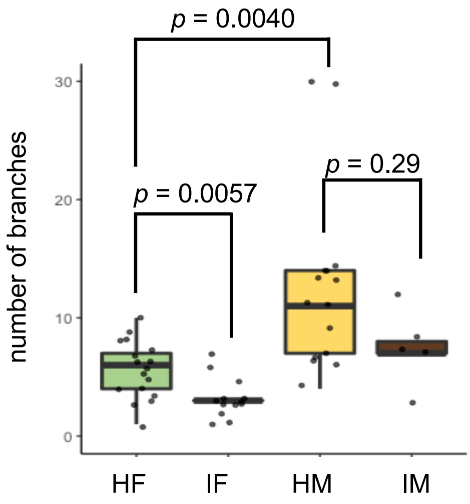


**Extended Data Fig.3: Effects of the anther smut fungus infection on flower numbers**

The flower number abundance in an individual was evaluated two weeks after the first flower opened. Genetically male plants (HM) produced more flowers than genetically female plants (HF) ($p$ = 0.0040 < 0.005, Two-sided Student's $t$-test with $N$ > 10), suggesting a sexual dimorphism in the flower numbers. Flower number was reduced in the infected female (IF) plants compared with HF ($p$ = 0.0057, Two-sided Student's $t$-test with N > 10). Similarly, the flower number was reduced in the infected male (IM), though it was not significant ($p$ = 0.29, Two-sided Student's $t$-test with $N$ =5).

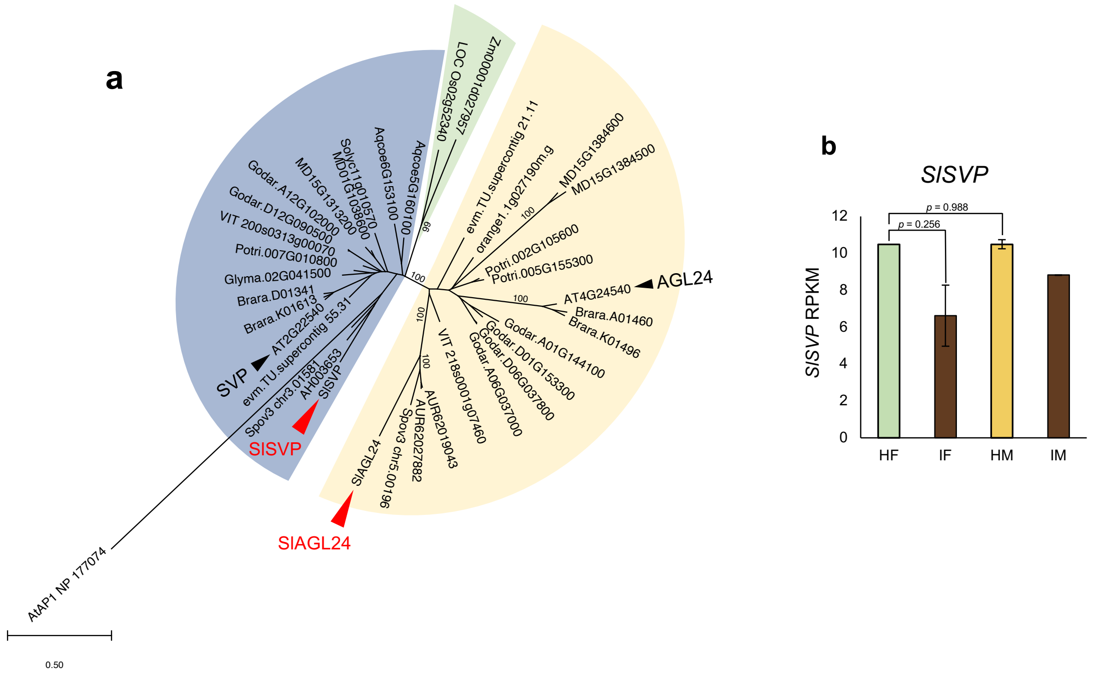


**Extended Data Fig.4: Phylogenetic analysis of *SlAGL24/SlSVP* and the expression levels of *SlSVP* in early inflorescence meristem**

**a,** Maximum likelihood-based phylogenetic analysis of *AGL24* and *SVP* orthologs identified in the whole genome sequences of 17 angiosperms, including *SlAGL24* and *SlSVP* sequences from *S. latifolia*, with *AtAP1* as the outgroup gene. *SlAGL24* and *A. thaliana AGL24* are nested into the clade (*yellow* clade) distinct from the clade including *SlSVP* and *A. thaliana SVP* (*blue* clade), with a statistic support (bootstrap = 100/100). **b,** The expression levels of *SlSVP* in early inflorescence meristems of genetically female (HF), the masculinized infected female (IF), genetically male (HM), and the infected male (IM). In contrast to the expression levels of *SlAGL24* (Extended Data Fig.4), *SlSVP* exhibited no substantial differences depending on the sex expressions or anther smut infections ($p$ = 0.988, 0.256, respectively).

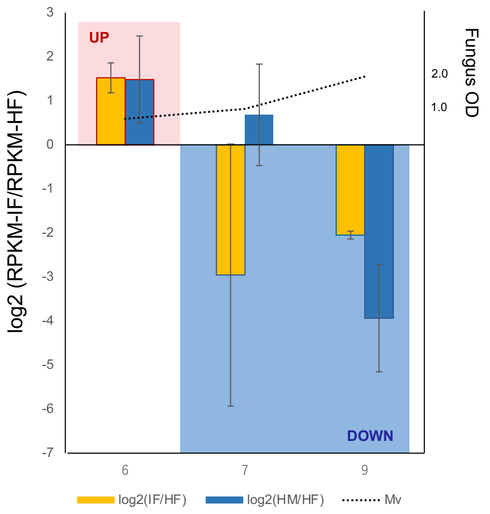


**Figure S5. Expression fluctuation of *SlAGL24*, in the suspension cell cultures of genetically male**

Transition of the *SlAGL24* expression patterns, according to *S. latifolia* suspension cell culturing (at day 6, 7, and 9). The expression bias between healthy female (HF) and anther smut fungus-infected female (IF), or $\log_2$(IF/ HF), and between healthy female (HF) and male (HM), or $\log_2$(HM/HF), were given in yellow and blue boxes, respectively. *SlAGL24* was upregulated until day 6, and subsequently downregulated after day 7 (see also Figure. 3I). The consistent trend was observed also in the comparison of healthy male (HM) and female (IF). The growth curve of the fungus ($OD_{600}$) was shown with the dashed line. *N* = 2 for each measuring point. Error bars show standard errors for each point.

**a**

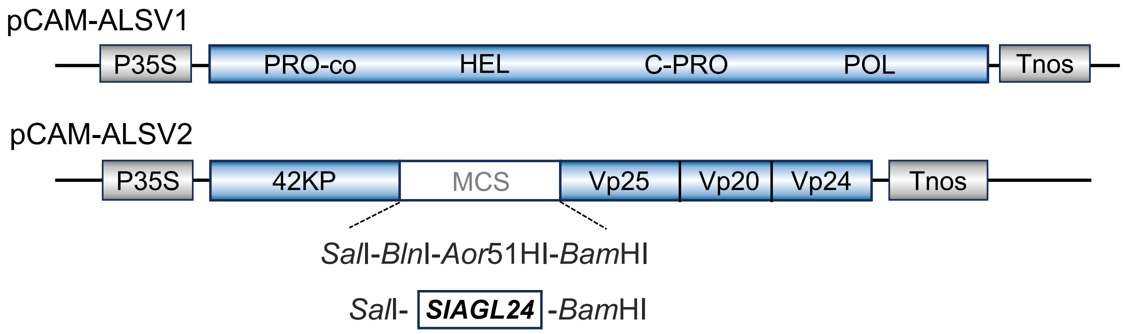


**b**

```
>SlAGL24_TRINITY_DN15866
ATGGCAAGGGAAAAGATAAAGATAAGGAAGATAGACAATATAACAGCAAGGCAAGTAACATTCTCAAAAAGAAGAAGA
GGGATTATTAAGAAAGCTGAAGAACTTGGTGTTCTTTGTGATGCCGATCTTGCCCTTATCATCTTTTCTTCTACTGGC
AAGGTCTTTGACTATTCCAGCTGCAGAATGAAGGATATCCTTACGCGTTACAAAATGCATTATTCGAAGAAGGAGACG
AGGCAAGAACAAACCTTCCTCGAGTTAAAGGAGCAAGAGGATAGCAACCTCGGTCGACTGAGAAAAGAGGCTCTGGAC
AAAAACAAGGAGCTTAGGAAATTTAGAGGGGAGGAACTTCATGGTCTAGACTTCAAGGAATTGCATGAACTCGAGCAA
ATGCTCGAGAACAGCTTATCTCGCGTTGTAGACATTAAGGAGAAGCGCTTAAGAAGTGAGATCGAGAATCTCCAAGAA
AAGGGTGAAATATTAATGGAAGAAAATACAAGGTTGAAACAAAGGTTGGCCATGTTATCACAAGGAAAGAAACCTGGA
AATTATGCTATCAAATCTGAGAGTAATACAACAGCAGAAGAGGGTCAATCATCTGAAACTGTTACTAATGGTGCTCCT
CCTCTCACTGATCAAGATGATAGTTCTGACACTTCTCTTAAATTAGGGTTAACCCTATTTTAA
```

**Figure S6. Construction of ALSV vector harboring a fragment of *SlAGL24* for VIGS in *S. latifolia* plant**

**a,** Schematic representation of the infectious cDNA clones of ALSV-RNA1 (pCAM-ALSV1) and ALSV-RNA2 (pCAM-ALSV2). **b,** A fragment of *SlAGL24* target gene (shown in white characters on a black background) was inserted between 42KP and VP25 using *Sal*I and *BamH*I sites. P35S, enhanced CaMV 35S promoter; Tnos, nopaline synthase terminator; PRO-co, protease cofactor; HEL, NTP-binding helicase; C-PRO, cysteine protease, POL, RNA polymerase; MP, 42K movement protein; Vp25, Vp20, and Vp24, capsid proteins.
